# Potentiating α_2_ subunit containing perisomatic GABA_A_ receptors protects against seizures in a mouse model of Dravet Syndrome

**DOI:** 10.1101/452813

**Authors:** Toshihiro Nomura, Nicole A. Hawkins, Jennifer A. Kearney, Alfred L. George, Anis Contractor

## Abstract

GABA_A_ receptor potentiators are commonly used for the treatment of epilepsy, but it is not clear whether distinct GABA_A_ receptor subtypes contribute to seizure activity, and whether targeting receptor subtypes will have disproportionate benefit over adverse effects. Here we demonstrate that the α_2_ / α_3_ selective positive allosteric modulator (PAM) AZD7325 preferentially potentiates hippocampal inhibitory responses at synapses proximal to the soma of CA1 neurons. The effect of AZD7325 on synaptic responses was more prominent in mice on the 129S6/SvEvTac background strain that has been demonstrated to be seizure resistant in the model of Dravet syndrome (*Scn1a*^+/−^) and in which the α_2_ GABA_A_ receptor subunits are higher relative to in the C57BL/6J strain. Consistent with this, treatment of mice with AZD7325 is associated with a higher temperature threshold for hyperthermia-induced seizures in *Scn1a*^+/−^ mice without apparent sedative effects. Our results in a model system indicate that selective targeting α_2_ is a potential therapeutic option for Dravet syndrome.

## Significance Statement

The GABA_A_ receptor is a target of several antiepileptic drugs (AEDs), but whether specific subtypes of GABA_A_ receptors are critical for seizure suppression is not known. We demonstrated that the α_2_ / α_3_ selective positive allosteric modulator (PAM) AZD7325 potentiates hippocampal inhibitory synaptic responses in mice in a strain-dependent manner. The results are consistent with our previous findings that uncovered strain dependent differential expression of the α_2_ subunit in seizure resistant and protective strains for *Scn1a* heterozygous deletion, a mouse model of Dravet syndrome. Treatment of mice with AZD7325 was associated with a higher temperature threshold for hyperthermia-induced seizures in *Scn1a*^+/−^ mice indicating α_2_ is a potential therapeutic target in Dravet syndrome.

## Materials & Methods

### Animals

All experimental procedures were carried out in accordance with the policies and protocols approved by the Northwestern University IACUC. Three separate mouse strains were used in the series of experiments; C57BL/6J (Strain # 000664, The Jackson Laboratory), 129S6/SvEvTac (Taconic), and [C57BL/6J x 129S6/SvEvTac] F1 strains. Male *Scn1a*^*tm1Kea*^ (*Scn1a*^+/−^) mice on the 129S6/SvEvTac background (129.*Scn1a*^+/−^)(Miller et al., 2014) were crossed with inbred female C57BL/6J or 129S6/SvEvTac mice to generate *Scn1a*^+/−^ mice with F1 (F1.*Scn1a*^+/−^) or 129S6/SvEvTac (129.*Scn1a*^+/−^) backgrounds, respectively. Both male and female *Scn1a*^+/−^ or wild-type littermates (*Scn1a*^+/+^) were used for experiments.

### Slice preparation for electrophysiology

Horizontal hippocampal slices (350 μm) were prepared from postnatal day 14 – 16 (P14 – P16) mice as described previously (Fernandes et al., 2015; Nomura et al., 2016). Briefly, brain sections were prepared in ice-cold sucrose-slicing artificial CSF (ACSF) containing the following (in mM): 85 NaCl, 2.5 KCl, 1.25 NaH_2_PO_4_, 25 NaHCO_3_, 25 glucose, 75 sucrose, 0.5 CaCl_2_, and 4 MgCl_2_ with 10 μM DL-APV and 100 μM kynurenate. Slices were incubated in a recovery chamber containing the same sucrose ACSF for ∼30 min at 30 °C, then the slicing solution was gradually exchanged for a recovery ACSF containing the following (in mM): 125 NaCl, 2.4 KCl, 1.2 NaH_2_PO_4_, 25 NaHCO_3_, 25 glucose, 1 CaCl_2_, and 2 MgCl_2_ with 10 μM DL-APV and 100 μM kynurenate at room temperature. After a recovery period of at least 1.5 h, slices were transferred to a recording chamber and visualized. During recordings, slices were continuously perfused with normal ACSF containing the following (in mM): 125 NaCl, 2.4 KCl, 1.2 NaH_2_PO_4_, 25 NaHCO_3_, 25 glucose, 2 CaCl_2_, and 1 MgCl_2_. Solution were equilibrated with 95 % O_2_ and 5 % CO_2_ throughout the experiment.

### Electrophysiological recording

Patch-clamp recordings were made from visually identified CA1 pyramidal neurons in the hippocampus. Recording electrodes had tip resistances of 3 – 5 MΩ when filled with internal recording solution containing the following (in mM): 135 CsCl, 20 HEPES, 2 EGTA, 2 Mg-ATP, 0.5 Na-GTP, and 10 QX-314 (pH 7.25). Monopolar extracellular stimulating electrodes were filled with ACSF and placed in the border of stratum radiatum (SR) and lacnosum moleculare (LM) or in stratum pyramidale (SP) to evoke distal or perisomatic synaptic responses, respectively (Prenosil et al., 2006; Jurgensen and Castillo, 2015). After forming the whole cell configuration, CA1 pyramidal neurons were voltage clamped at −70 mV. Inhibitory postsynaptic currents (IPSCs) were isolated by the inclusion of CNQX (10 μM) and D-APV (50 μM) in the extracellular solution. Miniature IPSCs (mIPSCs) were recorded in the presence of tetrodotoxin (TTX) (1 μM). To record asynchronous IPSCs (aIPSCs), the normal recording ACSF was exchanged for strontium-ACSF containing the following (in mM): 125 NaCl, 2.4 KCl, 1.2 NaH_2_PO_4_, 25 NaHCO_3_, 25 glucose, 6 SrCl_2_, 0.5 CaCl_2_, and 2 MgCl_2_. Stimuli were given through the extracellular stimulating electrodes placed in SP. Recorded aIPSCs were analyzed within a window from 50 ms to 500 ms after the stimulus artifact (Fernandes et al., 2015). Rise time was measured as the time from the onset of the inward current to the peak. Decay time was defined as the time from the peak to 37 % of the amplitude of the responses. Recordings were discarded when the access resistance (R_a_) showed > 20 % change during the recording. Data were collected and analyzed using pClamp 10 software (Molecular Devices) and Mini Analysis software (Synaptosoft).

### *In vivo* pharmacology and behavioral analysis

Male and female P18 - P20 F1.*Scn1a*^+/−^ mice were orally administered (gavage) 10, 17.8 or 31.6 mg/kg AZD7325 (generously provided by AstraZeneca, Inc.) or vehicle (5% 2-hydroxypropyl-β-cyclodextrin (HPBCD) (Acros Organics, New Jersey, USA)) 30 minutes prior to the induction of hyperthermia. Core body temperature was monitored using a RET-3 rectal temperature probe (Physitemp Instruments, Inc, New Jersey, USA) and controlled by a heat lamp connected to a rodent temperature regulator (TCAT-2DF, Physitemp) reconfigured with a Partlow 1160+ controller (West Control Solutions, Brighton, UK) as described previously (Hawkins et al., 2017). Body temperature was raised by 0.5 °C every two minutes until the onset of the first clonic convulsion with loss of posture. Once body temperature reached 42.5°C, the heat lamp was turned off and mice remained in the chamber until a generalized tonic-clonic seizure (GTCS) occurred or 3 minutes elapsed. Mice that did not experience a GTCS during the 3-minute hold were considered seizure-free.

Sedative effects of AZD7325 were assessed in the open field test. Wild-type P18 - P20 F1.*Scn1a*^+/+^ mice were administered 31.6 mg/kg AZD7325 or vehicle (5% HPBCD) by oral gavage 30 minutes prior to the analysis. Mice were then transferred to a 27.5 cm x 27.5 cm square acryl open field chamber and monitored for 60 min. Total distance traveled and the number of crossings into, and time mice spent in, the central zone (16.5 cm x 16.5 cm) were measured. Data were collected and analyzed using Ethovision 14 software (Noldus Information Technology).

### Data analysis

Statistical analyses were performed using GraphPad Prism and Origin Pro 9.0 software. Comparisons were made using Mann-Whitney *U* test for two samples. For comparisons for paired samples, Wilcoxon signed-rank test was employed. For multiple comparisons, unrepeated one-way or repeated two-way ANOVA followed by post hoc Bonferroni’s correction was used. For hyperthermia induced seizure threshold comparisons, time-to-event analysis was used with *p*-values determined by LogRank Mantel-Cox test. Differences were defined significant when *p* < 0.05. Data are presented as mean ± SEM.

## Introduction

Despite advances in the development of therapies for epilepsy, approximately 40% of patients have seizures that are refractory to treatment and also suffer from various comorbidities and adverse effects of antiepileptic drugs (AEDs) (Perucca et al., 2009; Brodie, 2017; Manford, 2017). The GABA_A_ receptor is a target of commonly used AEDs including benzodiazepines (BDZs), barbiturates, and valproate (VPA) (Greenfield, 2013). GABA_A_ receptors form heteromeric pentamers most typically with a stoichiometry composed by two α, two β, and one γ subunits (Olsen, 2018). The α_1_ and α_2_ subunits are amongst the most prevalent subunits present in synaptic GABA_A_ receptors, which participate in rapid phasic inhibition (Olsen and Sieghart, 2008).

The biological roles of α_1_ and α_2_ subunits are not fully characterized, however they govern the kinetics of the channel properties of GABA_A_ receptors. Receptors comprised of the α_2_ subunit mediate relatively slower decaying inhibitory postsynaptic currents (IPSCs) than those mediated by the most abundant GABA_A_ receptor type which contains α_1_ (α_1_β_2_γ_2_) (Gingrich et al., 1995) (Whiting, 2003). *In vivo*, activation of α_1_ containing GABA_A_ receptors is implicated in the hypnotic and motor effects of GABA receptor positive allosteric modulators (PAMs), whereas α_2_ and α_3_ containing receptors have been associated with anxiolytic and pain relieving effects (Low et al., 2000; McKernan et al., 2000; Knabl et al., 2008). The differential roles of these different receptor types in potentially controlling seizures is not clear, however subunit specific compounds hold promise for the treatment of epilepsies (Whiting, 2003; Rudolph and Knoflach, 2011).

Dravet Syndrome (DS) is an early onset epileptic encephalopathy (Scheffer et al., 2017; Steel et al., 2017) caused by haploinsufficiency of *SCN1A* that encodes the voltage-gated sodium channel Na_v_1.1 (Brunklaus and Zuberi, 2014; Steel et al., 2017). As with other monogenic disorders there is considerable evidence that pleiotropic effects of loss of *SCN1A* gene expression contribute to the phenotype. Particularly, work in a mouse model of DS (*Scn1a*^+/−^) has demonstrated that the cellular excitability and epilepsy phenotypes are highly dependent upon the background strain of the mouse (Yu et al., 2006; Miller et al., 2014; Mistry et al., 2014; Rubinstein et al., 2015; Hawkins et al., 2016), which may be a corollary to the variable clinical severity reported in patients (Mullen and Scheffer, 2009).

*Scn1a*^+/−^ mice on a congenic C57BL/6J background (B6.*Scn1a*^+/−^) or on a [C57BL/6J x 129S6/SvEvTac] F1 hybrid background (F1.*Scn1a*^+/−^) develop a severe phenotype with spontaneous seizures and premature lethality. In contrast, *Scn1a*^+/−^ mice on a congenic 129S6/SvEvTac background (129.*Scn1a*^+/−^) do not develop spontaneous seizures and have longevity comparable to wild-type littermates (Miller et al., 2014). These strain-dependent phenotypes have led to the suggestion that modifier genes strongly contribute to the phenotype severity (Mullen and Scheffer, 2009; Miller et al., 2014; Mistry et al., 2014; Rubinstein et al., 2015). Genetic mapping studies have discovered several candidate genes contributing to the strain-dependent phenotype in mice (Miller et al., 2014; Hawkins and Kearney, 2016; Hawkins et al., 2016; Calhoun et al., 2017).

One of the candidate modifier genes, *Gabra2*, is of particular interest because it encodes for the α_2_ subunit of the GABA_A_ receptor (Hawkins et al., 2016). These studies demonstrated higher *Gabra2* transcript and protein expression in the 129 protective strain compared to expression in the two strains that are susceptible to the *Scn1a* mutation, B6 and F1. This strain-dependence of expression of GABA receptor subunits is consistent with previous reports (Mulligan et al., 2012; Yeo et al., 2016). Moreover, the administration of the 1,5-BDZ clobazam (CLB) is associated with higher temperature threshold for hyperthermia-induced seizure in F1.*Scn1a*^+/−^ mice, which indicates that potentiating GABA_A_ receptors has a strong anti-seizure effect in these mice (Hawkins et al., 2016; Hawkins et al., 2017). This is consistent with CLB and VPA as first-line treatments in DS patients (Wirrell et al., 2017).

Here we demonstrated that AZD7325, a selective PAM for α_2_ / α_3_ subunit containing GABA_A_ receptors, preferentially potentiated GABAergic inhibitory postsynaptic currents (IPSCs) in hippocampal perisomatic synapses by prolonging their decay kinetics. There was a strong strain dependent effect of the drug, with a significantly larger effect in the 129 mouse strain than in B6 or F1 strains. These findings are consistent with the interpretation that α_2_ GABA receptor subunits are relatively enriched in synapses of mice in the 129 strain. In addition, *in vivo* administration of AZD7325 was associated with a higher temperature threshold for hyperthermia-induced seizure in *Scn1a*^+/−^ mice without any apparent sedative effect. Our results indicate that the α_2_ subunit is critical for seizure pathophysiology in the DS mouse model and targeting α_2_ containing receptors with a subunit selective PAM is a potential therapeutic strategy for DS.

## Results

### AZD7325 is a PAM for native GABA_A_ receptors

AZD7325, 4-amino-8-(2-fluoro-6-methoxy-phenyl)-N-propyl-cinnoline-3-carboxamide, has been shown to have selectivity for α_2_ / α_3_ subunits and to potentiate GABA_A_ receptor mediated responses in heterologous expression systems (Alhambra et al., 2011; Saito et al., 2016). To determine how AZD7325 affects native GABA receptors, we examined the effect of the compound on phasic inhibitory synaptic events in hippocampal slices of wild-type F1 mice. Miniature IPSCs (mIPSCs) were recorded from CA1 pyramidal neurons, before and after the application of vehicle (Veh; DMSO) or AZD7325 (AZD; 100 nM) for ∼10 min. AZD7325 treatment did not potentiate the amplitude (Figure 1 a and b) but significantly prolonged the decay time of mIPSCs (Figure 1 c – e). There was no effect on the frequency and the rise time of mIPSCs after the AZD7325 treatment (Wilcoxon signed-rank test: Frequency: 4.3 ± 0.5 Hz and 4.2 ± 0.4 Hz, *p* = 0.41; Rise time: 2.4 ± 0.1 ms and 2.5 ± 0.1 ms, *p* = 0.33; before and after the treatment, respectively; n = 15 cells / 7 mice). These results demonstrate that AZD7325 acts as a PAM for native GABA_A_ receptors by prolonging the decay kinetics of IPSCs in the hippocampus in acute slices.

**Figure 1:**
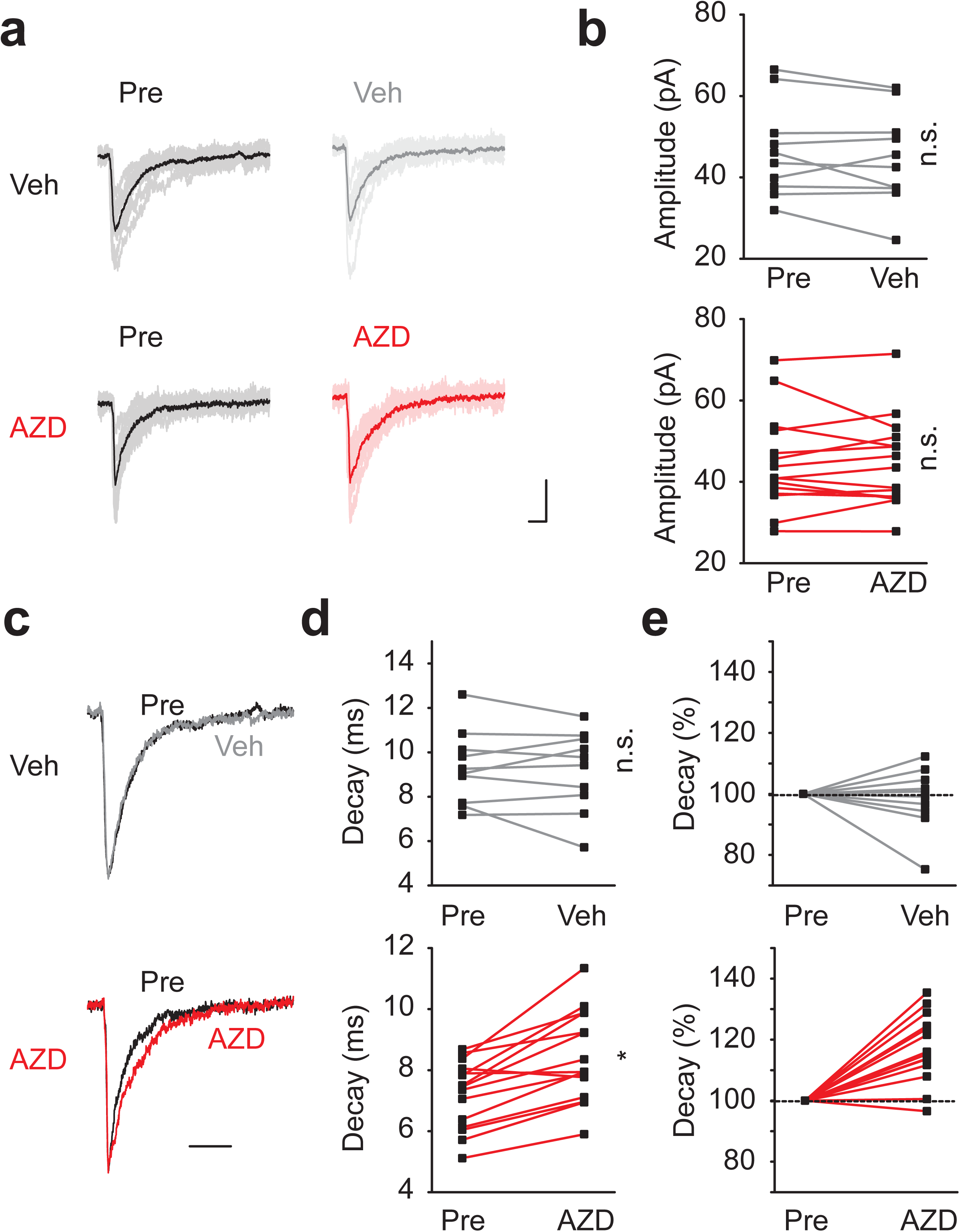
**AZD7325 potentiated Inhibitory synaptic currents in CA1 neurons** (**a**) Representative mIPSC traces before (Pre) and after DMSO (Veh; top) or AZD7325 (AZD; bottom) treatments. 10 representative IPSCs and the averaged trace are presented in thin and thick lines, respectively. Calibration: 10 ms, 30 pA. (**b**) Grouped data from all recordings of mIPSC amplitude before (Pre) and after DMSO (Veh) or AZD7325 (AZD) treatments. Amplitudes are not affected by the treatment. Wilcoxon signed-rank test: Veh group: 46.5 ± 3.6 pA (Pre) and 44.7 ± 3.7 pA (Veh), p = 0.28, n = 10 cells / 5 mice; AZD group: 44.6 ± 3.0 pA (Pre) and 44.5 ± 2.8 pA (AZD), p = 0.72, n = 15 cells / 7 mice. n.s. denotes p ≥ 0.05. (**c**) Representative mIPSC traces scaled to the peak before (Pre) and after DMSO (Veh) or AZD7325 (AZD) treatments. Calibration: 20 ms. (**d**) Grouped data for mIPSC decay time before (Pre) and after DMSO (Veh) or AZD7325 (AZD) treatments. Decay is prolonged after AZD treatment. Wilcoxon signed-rank test: Veh group: 9.3 ± 0.5 ms (Pre) and 9.2 ± 0.6 ms (Veh), p = 0.85, n = 10 cells / 5 mice; AZD group : 7.2 ± 0.3 ms (Pre) and 8.4 ± 0.4 ms (AZD), p < 0.01, n = 15 cells / 7 mice. n.s. and * denotes p ≥ 0.05 and < 0.05, respectively. (**e**) Relative mIPSC decay time before (Pre) and after DMSO (Veh) or AZD7325 (AZD) treatments.

### AZD7325 has differential activity at perisomatic synapses

AZD7325 has been reported to have high affinity for α_2_ and α_3_ subunits of GABA_A_ receptors (Alhambra et al., 2011; Saito et al., 2016). In the hippocampus, prior work has demonstrated that GABA_A_ receptor content of inhibitory synapses is spatially distributed along the somato-dendritic length. Synapses close to the soma preferentially contain α_2_ GABA_A_ receptor subunits whereas synapses more distal from the soma are dominated by α_1_ subunits (Prenosil et al., 2006). To differentially examine perisomatic and distal synapses, we isolated evoked IPSCs from either population by stimulating inhibitory synaptic input in different layers in the hippocampal slice (Prenosil et al., 2006; Jurgensen and Castillo, 2015) (Figure 2a). As expected, perisomatically evoked IPSCs had kinetics distinct from the distally evoked synaptic currents, which displayed significantly faster rise times and decay times (Figure 2 b and c). AZD7325 had no effect on the decay kinetics of distal IPSCs (Figure 2 d – f), whereas the decay kinetics of perisomatic evoked IPSCs were significantly prolonged by AZD7325 application (Figure 2 d – f). These results demonstrate that the α_2_ / α_3_ selective PAM differentially modulates inhibitory synapses located in the perisomatic region of CA1 neurons. This finding is consistent with the proposal that there are segregated populations of GABA_A_ receptors at CA1 synapses (Prenosil et al., 2006). As with mIPSCs, these were no significant effects of AZD7325 on the amplitude of IPSCs in either distal or perisomatic synapses (Wilcoxon signed-rank test: Distal: 216.0 ± 27.0 pA and 197.8 ± 31.1 pA, p = 0.23; Perisomatic: 1034.0 ± 118.6 pA and 913.4 ± 149.5 pA, p = 0.16; before and after the treatment, respectively: n = 10 cells / 5 mice and n = 7 cells / 4 mice, in distal and perisomatic synapses, respectively). In addition, the paired pulse ratio (PPR) measured at an interval of 50 ms was not affected by AZD7325 suggesting that this compound does not have any obvious unintended presynaptic effects (Figure 2 g & h).

**Figure 2:**
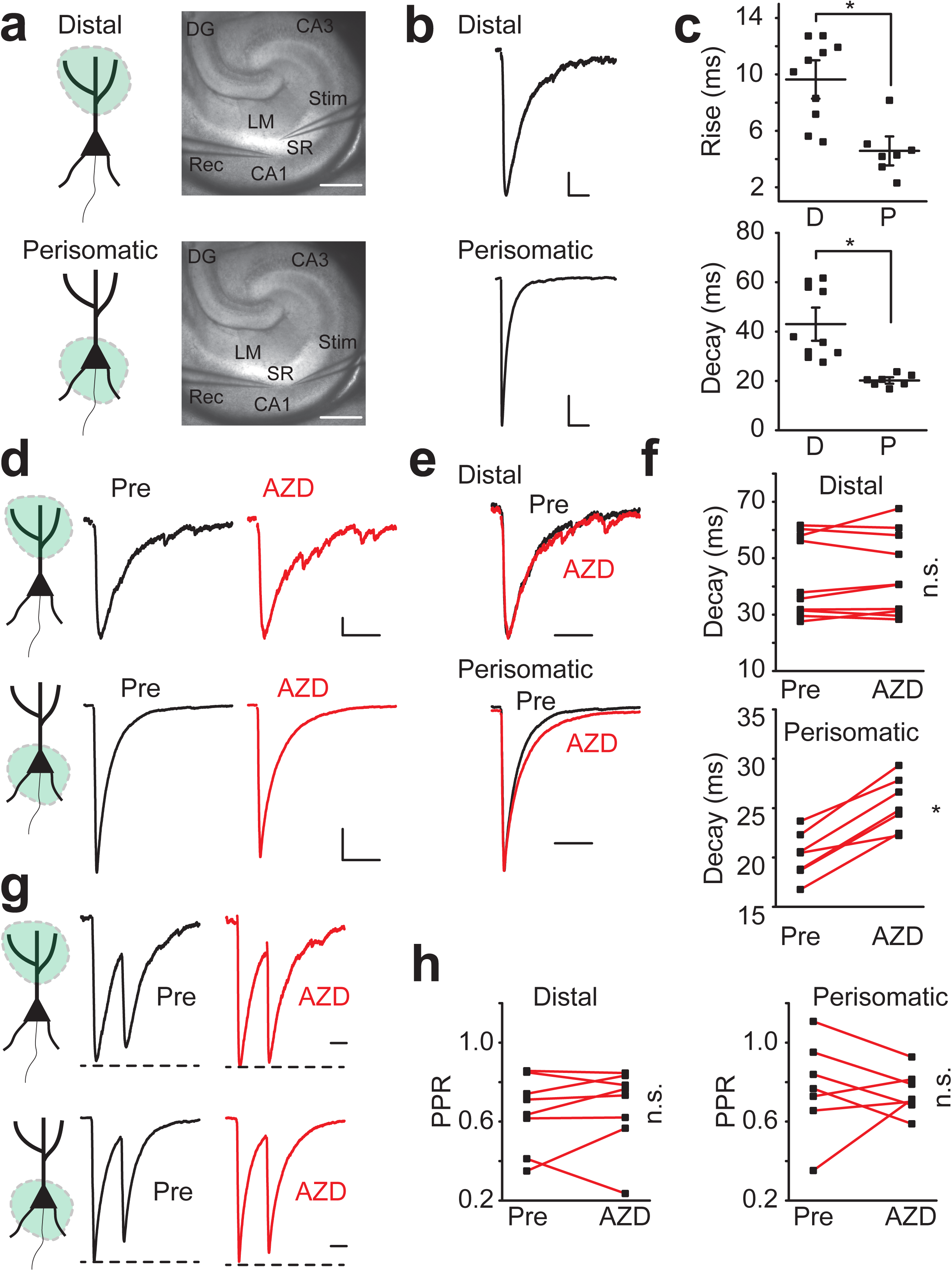
**AZD7325 potentiates perisomatic but not distal inhibitory inputs to CA1 pyramidal neurons** (**a**) Cartoon representation of the region activated, and image of the hippocampus with position of stimulating and recording electrodes for distal stimulation (top) and perisomatic stimulation (bottom) of IPSCs. Calibration: 500 μm, Abbreviations; SR : Stratum Radiatum, LM : Lacunosum Moleculare, CA1 : Cornus Ammonis 1, CA3 : Cornus Ammonis 3, DG : Dentate Gyrus, Rec : recording electrode, Stim : stimulating electrode. (**b**) Representative evoked IPSC traces in distal (top) and perisomatic (bottom) synapses. Calibration: 50 ms, 20 pA (distal), 200 pA (perisomatic). (**c**) Grouped data for all evoked IPSC recordings summarizing rise times and decay time in distal (D) and perisomatic (P) synapses. Mann-Whitney *U* test : Rise time: 9.6 ± 0.9 ms (D) and 4.6 ± 0.7 ms (P), p < 0.01; Decay time: 43.0 ± 4.5 ms (D) and 20.2 ± 0.9 ms (P), p < 0.01, n = 10 cells / 5 mice (D) and 7 cells / 4 mice (P) * denotes p < 0.05 (**d**) Representative IPSC traces in distal (top) and perisomatic (bottom) synapses before (Pre) and after AZD7325 (AZD) treatment. Calibration: 50 ms, 20 pA (distal), 200 pA (perisomatic). (**e**) Representative scaled IPSC traces evoked in distal (top) and perisomatic (bottom) synapses before (Pre) and after AZD7325 (AZD) treatment. Calibration: 50 ms (**f**) Grouped data from all recordings of IPSC decay time in distal and perisomatic synapses before (Pre) and after AZD7325 (AZD) treatment. IPSC decay is prolonged specifically in perisomatic synapses. Wilcoxon signed-rank test: Distal: 43.0 ± 4.5 ms (Pre) and 44.0 ± 4.6 ms (AZD), p = 0.63, n = 10 cells / 5 mice; Perisomatic: 20.2 ± 0.9 ms (Pre) and 25.4 ± 1.0 ms (AZD), p = 0.016, n = 7 / 4 mice. n.s. and * denotes p ≥ 0.05 and < 0.05, respectively. (**g**) Representative paired IPSC traces at 50 ms inter-stimulus interval in distal (top) and perisomatic (bottom) synapses before (Pre) and after AZD7325 (AZD) treatment. Calibration: 20 ms (**h**) Grouped data for all recordings of paired-pulse ratio (PPR) in distal and perisomatic synapses before (Pre) and after AZD7325 (AZD) treatment. PPR is not altered by AZD treatment. Wilcoxon signed-rank test: Distal: 0.65 ± 0.07 (Pre) and 0.67 ± 0.07 (AZD), p = 0.55, n = 8 cells / 5 mice; Perisomatic: 0.77 ± 0.09 ms (Pre) and 0.75 ± 0.04 ms (AZD), p = 0.58, n = 7 cells / 4 mice. n.s. denotes p ≥ 0.05.

### The magnitude of AZD7325 effect is dependent upon the background strain

We and others have previously reported that there is marked strain dependence in the expression levels of the α_2_ subunit of GABA_A_ receptors(Mulligan et al., 2012; Hawkins et al., 2016; Yeo et al., 2016). This strain dependent expression difference is correlated with phenotype severity in the DS mouse model, suggesting that it may contribute to the underlying protective effect from seizures in 129.*Scn1a*^+/−^ mice (Hawkins et al., 2016). We therefore tested whether AZD7325 has a strain dependent effect. AZD7325 significantly prolonged the decay time of mIPSCs recorded from CA1 neurons from mice on the 129 background strain (Figure 3 a – f). The decay time was lengthened by 130.0 ± 3.2 % (n = 13) after AZD7325 treatment in the 129 strain, which was significantly greater than that in B6 (114.4 ± 2.9 %, n = 12) and F1 (117.1 ± 2.8 %, n = 15 : Figure 1 e) (one-way ANOVA : *F*_(2,37)_ = 7.69, *p* = 0.003 for B6 vs 129, 0.01 for F1 vs 129) mice (Figure 3 f; F1 data not shown in the figure). These results are consistent with our previous report of differential expression of *Gabra2* gene in the 129 strain that confers resistance to seizures in the *Scn1a*^+/−^ DS model (Hawkins et al., 2016).

**Figure 3:**
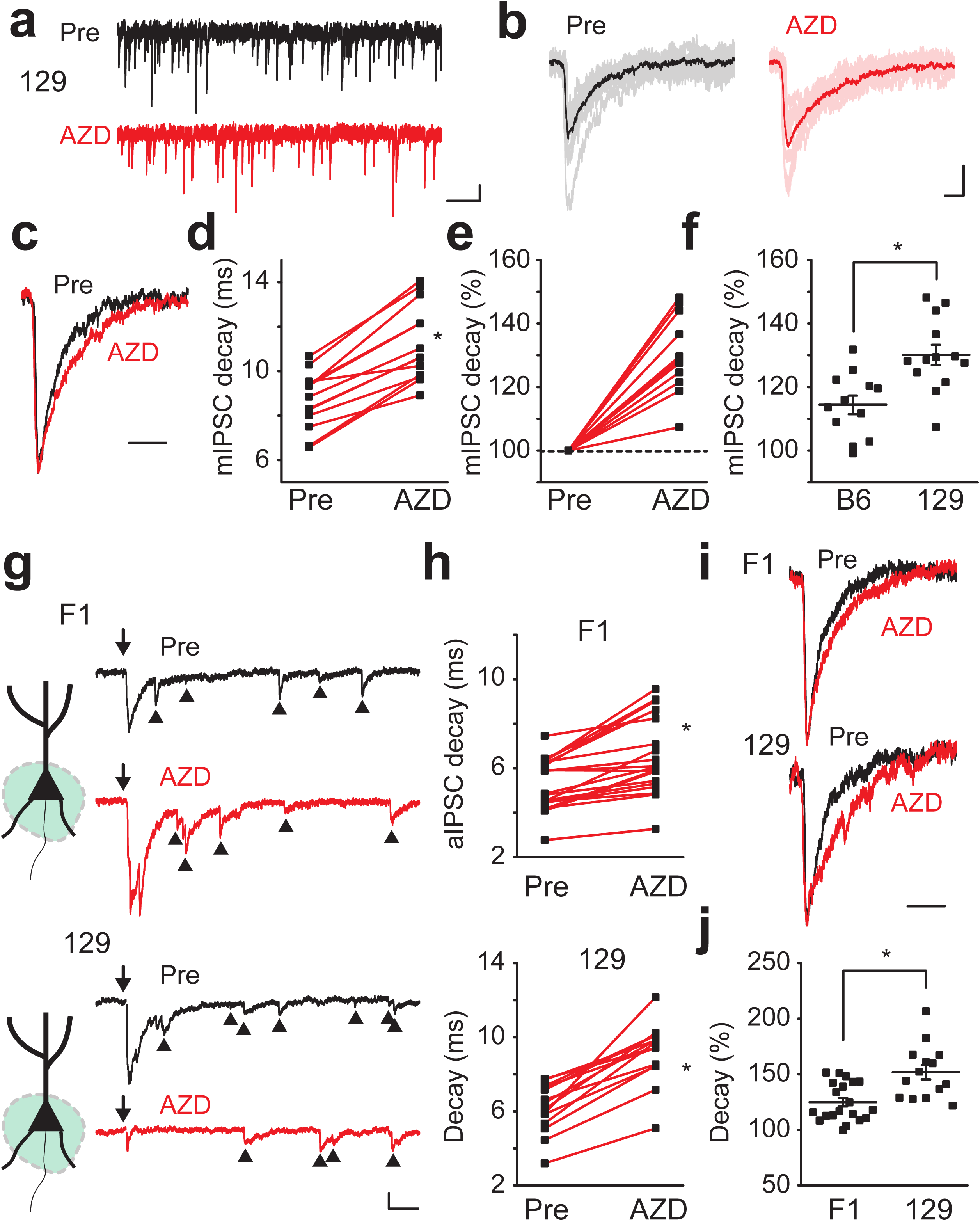
**Strain dependent effect of AZD7325 on IPSCs** (**a**) Representative 10 second mIPSC traces before (Pre) and after AZD7325 (AZD) treatment in 129 mice. Calibration: 1 s, 20 pA. (**b**) Representative individual mIPSC traces. 10 representative and averaged traces were presented in thin and thick lines, respectively. Calibration: 10 ms, 20 pA. (**c**) Representative scaled mIPSC traces before (Pre) and after AZD7325 (AZD) treatment. Calibration: 20 ms. (**d**) Grouped data for mIPSC decay time before (Pre) and after AZD7325 (AZD) treatment. Wilcoxon signed-rank test: Mean decay time: 8.7 ± 0.4 ms (Pre) and 11.2 ± 0.5 ms (AZD), p < 0.01, n = 13 cells / 6 mice * denotes p < 0.05. (**e**) Relative mIPSC decay time before (Pre) and after AZD7325 (AZD) treatment. (**f**) Grouped data for mean mIPSC decay times after AZD7325 (AZD) treatment in C57BL/6J (B6) and 129S6/SvEvTac (129) strains. AZD induced decay prolongation is greater in 129 strain than in B6 strain. One-way ANOVA: Relative decay time: 114.4 ± 2.9 % (B6) and 130.1 ± 3.2 % (129), p < 0.01, n = 12 cells / 3 mice (B6) and 13 cells / 6 mice (129). * denotes p < 0.05. (**g**) Representative aIPSC traces (550 ms) before (Pre) and after AZD7325 (AZD) treatments in F1 and 129 strains. Arrows (↓) indicate time at which inputs were stimulated. Arrow heads (▴) denote asynchronous events. Calibration: 50 ms, 50 pA. (**h**) Grouped data for all recordings of aIPSC decay time before (Pre) and after AZD7325 (AZD) treatment in F1 and 129 strains. aIPSC decay time is prolonged by AZD treatment in both strains. Wilcoxon signed-rank test: F1 strain: 5.3 ± 0.3 ms (Pre) and 6.6 ± 0.4 ms (AZD), p < 0.01, n = 20 cells / 5 mice; 129 strain: 6.2 ± 0.4 ms (Pre) and 9.2 ± 0.4 ms (AZD), p < 0.01, n = 14 cells / 6 mice * denotes p < 0.05. (**i**) Representative scaled aIPSC traces before (pre) and after AZD7325 (AZD) treatment in F1 and 129 strains. Calibration: 20 ms. (**j**) Grouped data for modulation of aIPSC decay time by AZD7325 (AZD) in F1 and 129 strains. AZD induced decay prolongation is greater in 129 strain than in F1 strain. Mann-Whitney *U* test: Relative decay time: 124.9 ± 3.9 % (F1) and 151.8 ± 6.4 % (129), p < 0.01, n = 20 cells / 5 mice (F1) and 14 cells / 6 mice (129). * denotes p < 0.05.

Because our local stimulation experiments had demonstrated that AZD7325 is effective only on perisomatic stimulated synapses, we selectively stimulated these whilst also using strontium (Sr^2+^) replacement to isolate individual synaptic events by desynchronizing release of the compound IPSCs and to record asynchronous IPSCs (aIPSCs) in the perisomatic synapses (Xu-Friedman and Regehr, 2000; Fernandes et al., 2015). AZD7325 application significantly prolonged the decay time of aIPSCs in the perisomatic synapses in both F1 and 129 strains. However, the effect of AZD7325 on the decay time of aIPSCs was greater in the 129 strain compared to the F1 strain, consistent with a differential expression of the α_2_ subunit in inhibitory synapses in these strains (Figure 3 g – j).

### AZD7325 modulates IPSCs in *Scn1a*^+/−^ mice

Our data indicate that AZD7325 specifically modulates inhibitory currents particularly in perisomatic synapses in the CA1 of the hippocampus and suggest that potentiation of α_2_ subunit containing GABA_A_ receptors could be seizure protective in *Scn1a*^+/−^ mice. Therefore, we tested whether AZD7325 modulates hippocampal GABAergic responses in *Scn1a*^+/−^ mice. We first examined the effect of AZD7325 on hippocampal inhibitory synaptic transmission in CA1 neurons in *Scn1a*^+/−^ mice. AZD7325 application prolonged the decay time of mIPSCs in *Scn1a*^+/−^ mice, and as expected the modulation was greater in 129.*Scn1a*^+/−^ mice on the seizure resistant strain than in the seizure susceptible F1.*Scn1a*^+/−^ strain (Figure 4 a – c). Similarly, AZD7325 significantly prolonged the aIPSC decay time in perisomatic synapses, and consistent with the mIPSC findings, the prolongation of the decay kinetics was stronger in slices from mice on the 129 background (Figure 4 d – f).

**Figure 4:**
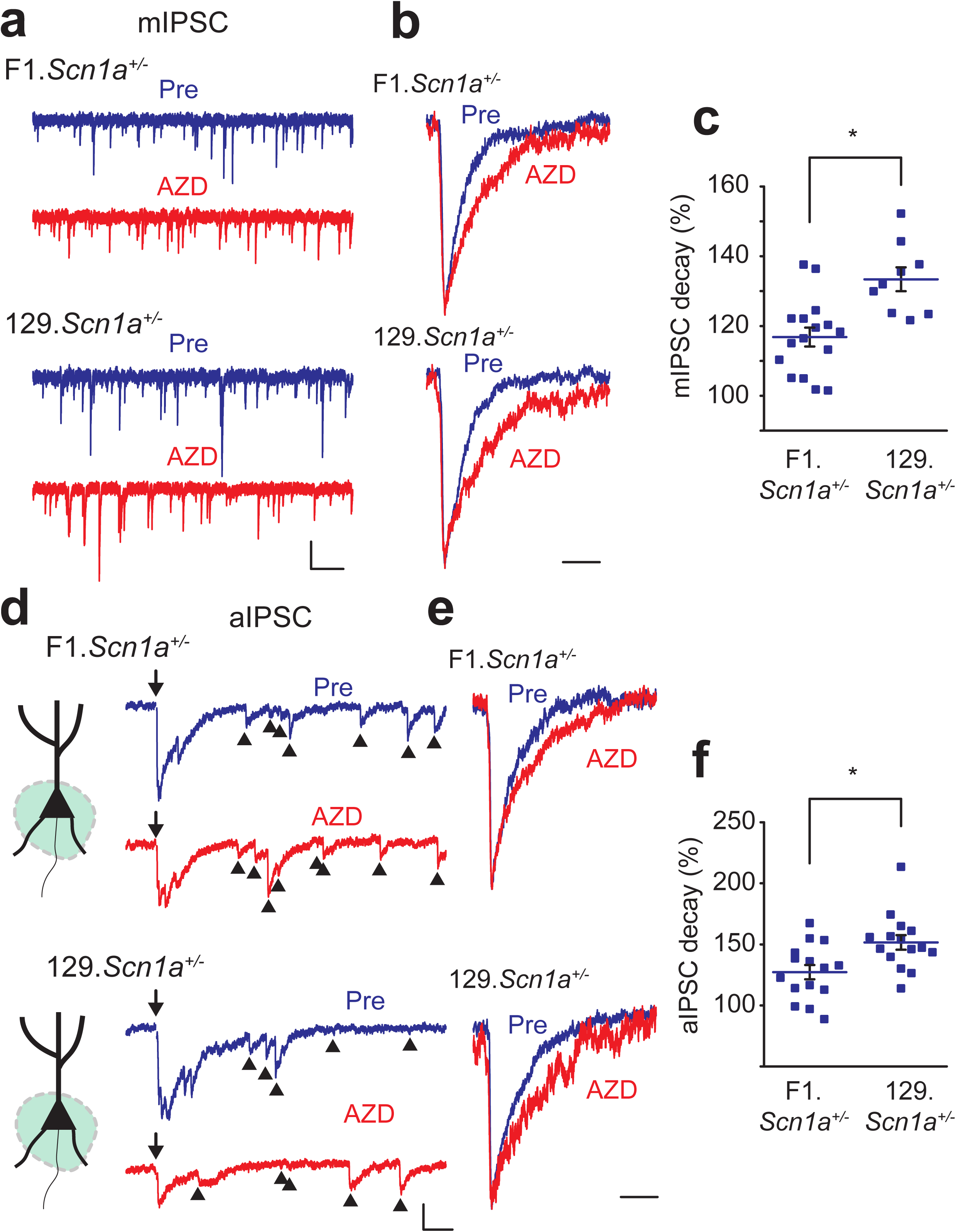
**AZD7325 is more effective in modulating IPSCs in 129.*Scn1a*^+/−^ mice than F1.*Scn1a*^+/−^ mice**. (**a**) Representative mIPSC traces (10 s) before (Pre) and after AZD7325 (AZD) treatments in F1.*Scn1a*^+/−^ and 129.*Scn1a*^+/−^ mice. Calibration: 1 s, 40 pA. (**b**) Scaled individual mIPSC traces before (Pre) and after AZD7325 (AZD) treatments in F1.*Scn1a*^+/−^ and 129.*Scn1a*^+/−^ mice. Calibration: 20 ms. (**c**) Grouped data for all mIPSC in F1.*Scn1a*^+/−^ and 129.*Scn1a*^+/−^ mice. Mann-Whitney *U* test: Mean relative decay times after AZD7325 treatment: 116.8 ± 2.7 % (F1.*Scn1a*^+/−^) and 133.4 ± 3.4 % (129.*Scn1a*^+/−^), p < 0.01, n = 16 cells / 6 mice and 9 cells / 4 mice. * denotes p < 0.05. (**d**) Representative aIPSC traces before (pre) and after AZD7325 (AZD) treatments in F1.*Scn1a*^+/−^ and 129.*Scn1a*^+/−^ mice. Arrows (↓) indicate time at which inputs were stimulated. Arrow heads (▴) denote asynchronous events. Calibration: 50 ms, 50 pA. (**e**) Scaled aIPSC traces before (pre) and after AZD7325 (AZD) treatments. Calibration: 20 ms. (**f**) Grouped data for all aIPSC in F1.*Scn1a*^+/−^ and 129.*Scn1a*^+/−^ mice. AZD effect is greater in 129.*Scn1a*^+/−^ than in F1.*Scn1a*^+/−^ mice. Mann-Whitney *U* test: Mean relative decay times after AZD7325 treatments: 127.2 ± 5.9 % (F1.*Scn1a*^+/−^) and 151.6 ± 6.0 % (129.*Scn1a*^+/−^), p < 0.01, n = 15 cells / 4 mice and 15 cells / 6 mice. * denotes p < 0.05.

### AZD7325 alters the threshold for hyperthermia - induced seizures in *Scn1a*^+/−^ mice

To directly assess whether *in vivo* AZD7325 administration affects the seizure threshold in F1.*Scn1a*^+/−^ mice, we used a hyperthermia-induced seizure assay (Hawkins et al., 2016). AZD7325 had a significant protective effect against seizure induction in F1.*Scn1a*^+/−^ mice (Figure 5 a). At doses as low as 10 mg/kg, there was a significantly higher threshold temperature for hyperthermia-induced seizures. At higher doses up to 31.6 mg/kg, this effect was sustained but not significantly greater than lower doses (Figure 5 a). Importantly, no obvious signs of sedation were noted during the hyperthermia assay. To quantify whether the highest dose of AZD7325 was sedating, we tested another cohort of wild-type F1 mice and monitored their activity in an open field assay. Both vehicle and AZD7325 treated mice demonstrated a characteristic habituation of activity during the 60 min monitoring. However, no difference was observed in the total activity or distance traveled between mice in the vehicle and AZD7325 treatment groups, demonstrating no major sedation caused by AZD7325 at this dose (Figure 5 b – d). The time spent in the center of the chamber and the number of crossings into the center were indistinguishable between the groups, suggesting AZD7325 did not show an obvious anxiolytic effect in wild-type mice at this dose (Figure 5 e – f).

**Figure 5:**
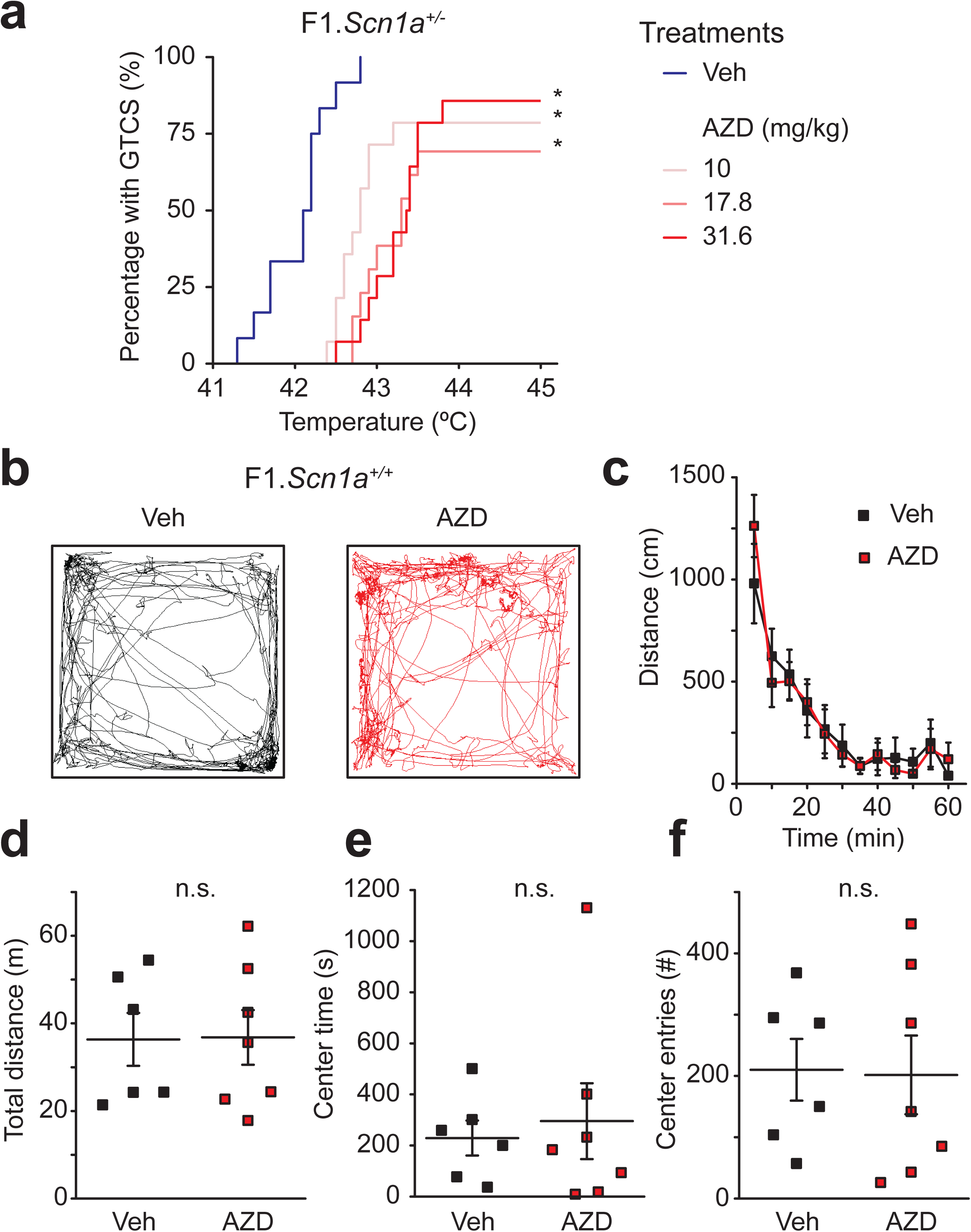
**AZD7325 treatment attenuates hyperthermia-induced seizures in F1.*Scn1a*^+/−^ mice with no sedative effect.** (**a**) Cumulative generalized tonic-clonic seizure (GTCS) incidence curve for F1.*Scn1a*^+/−^ mice treated with AZD7325 (AZD) or vehicle (Veh) and then subjected to hyperthermia-induced seizure threshold testing. All three doses, 10, 17.8 and 31.6 mg/kg significantly shifted the temperature threshold curve for hyperthermia-induced seizures (p < 0.0001). Vehicle treated F1.*Scn1a*^+/−^ mice had median GTCS temperature threshold of 42.2°C, compared to median thresholds in the treatment groups of 42.8°C for 10 mg/kg, 43.3°C for 17.8 mg/kg, and 43.4°C for 31.6 mg/kg. *P*-values were determined by LogRank Mantel-Cox with n = 12 - 14 per treatment group. (**b**) Representative trajectory maps of the open field test in vehicle (Veh) or AZD7325 (AZD) administered mice. (**c**) Collective data for the time course of the distance travelled by vehicle (Veh) or AZD7325 (AZD) administered mice in the open field. Both groups show a time dependent decrease in the distance reflecting habituation to the new environment (two-way ANOVA : *F*_(11,121)_ = 22.95, *p* < 0.0001, 5 min bins, n = 6 and 7 mice for vehicle and AZD7325 treated groups, respectively), but there is no difference between the treatment groups (two-way ANOVA : *F*_(1,11)_ = 0.0030, *p* = 0.96, n = 6 and 7 mice for vehicle and AZD7325 treated groups, respectively). (**d**) Grouped data of effect of AZD on total distance travelled. AZD7325 treatment did not have an effect on total distance travelled. Mann-Whitney *U* test: Total distance: 36.3 ± 6.0 m (Veh) and 36.8 ± 6.2 m (AZD), p = 1.0, n = 6 and 7 mice for vehicle and AZD7325 treated groups. n.s. denotes p ≥ 0.05. (**e** & **f**) The time mice spent in the center (**e**) and the number of the times mice crossed into the center of the open field chamber (**f**) Mann-Whitney *U* test: Center time: 230 ± 68 s (Veh) and 295 ± 148 s (AZD), p = 0.73 (**e**) Center enter number: 210 ± 50 times (Veh) and 202 ± 64 s (AZD), p = 0.76 (**f**) n = 6 and 7 mice for vehicle and AZD7325 treated groups. n.s. denotes p ≥ 0.05.

## Discussion

In this study, we demonstrated that AZD7325, which has previously been characterized as a PAM selective for recombinant α_2_ / α_3_ subunits of GABA_A_ receptors (Alhambra et al., 2011), potentiates hippocampal GABAergic synaptic responses in the CA1, and is protective for hyperthermia induced seizures in the F1.*Scn1a*^+/−^ mice, a mouse model of DS. Moreover, we observed that GABAergic synapses proximal to the somatic region of CA1 neurons are preferentially modulated by AZD7325, suggesting that there is a distributed localization of α_2_ / α_3_ containing GABA_A_ receptors that might provide a potent inhibitory influence on pyramidal neurons. The results are the first to demonstrate the action of this compound on native receptors in the hippocampus and to provide insight into how it enhances phasic inhibitory transmission, which may contribute to the dampening of seizure activity.

### GABA_A_ receptors containing α_2_ subunits have a privileged role in hippocampal inhibition

GABA_A_ receptors are formed from heterogeneous assemblies of subunits that are both developmentally and regionally regulated to give diverse expression patterns throughout the brain (Mody and Pearce, 2004). The α subunits are of particular interest for targeted therapeutics because of the divergence in their activities that segregate the sedative and anesthetic effects mediated by α_1_ subunit containing GABA_A_ receptors (Whiting, 2003; Rudolph and Knoflach, 2011) from the anxiolytic and analgesic effects of drugs selectively targeting α_2_ / α_3_ subunit GABA_A_ receptors. Thus, it is possible that targeting subsets of GABA_A_ receptors containing the α_2_ / α_3_ subunits could be used to mitigate undesirable sedative and amnesic effects of GABAergic AEDs. Several PAMs selective for the α_2_ / α_3_ subunits have been developed including L-838,417 (α_1_-sparing PAM) (McKernan et al., 2000; Knabl et al., 2008), TCS1105 (Taliani et al., 2009), AZD7325 (Alhambra et al., 2011; Zhou et al., 2012), and PF-06372865 (Gurrell et al., 2018; Nickolls et al., 2018), although none of these are currently in clinical use as an AED. The commonly prescribed AED and anxiolytic medication clobazam (CLB) is a 1,5-BDZ that is known to be metabolized *in vivo* to active N-desmethyl clobazam (N-CLB) that has modest preference for α_2_ / α_3_ containing receptors (Ralvenius et al., 2016) (but, also see (Jensen et al., 2014)).

In direct recordings from CA1 neurons we observed that selective stimulation of perisomatic, but not distal dendritic IPSCs could isolate synapses that were preferentially potentiated by the α_2_ / α_3_ selective PAM AZD7325. This is consistent with previous reports that demontrated α_1_ containing receptors are functionally localized to distal dendritic synapses whereas α_2_ containing GABA_A_ receptors preferentially mediate perisomatic phasic inhibition (Prenosil et al., 2006). Using high resolution EM freeze fracture immunolabeling, it was recently demonstrated that synapses on the soma and axon initial segment (AIS) contain receptors with both α_1_ and α_2_ subunits (Kerti-Szigeti and Nusser, 2016). Moreover, prior work had suggested a remarkable input specific molecular segregation of GABA_A_ receptor subunits. Specifically, α_1_ containing receptors at synapses formed by paralbumin (PV) positive interneurons (Klausberger et al., 2002) and α_2_ containing receptors at synapses formed by cholecystokinin (CCK) positive interneurons (Nusser et al., 1996; Fritschy et al., 1998; Nyiri et al., 2001) may not be so clearly divided. Rather, distinct synapses onto the soma from PV, CCK, and those onto the AIS have similar subunit composition where both α_1_ and α_2_ subunits are present in most labeled synapses (Kasugai et al., 2010; Kerti-Szigeti and Nusser, 2016). Our functional results using the selective pharmacological profile of AZD7325 are consistent with inhibitory synapses localized close to the soma containing receptors with the α_2_ subunit.

### Modulation of GABA_A_ receptors containing α_2_ subunits suppresses seizures

GABA_A_ receptor PAMs are known to potentiate IPSCs by increasing the amplitude, as well as prolonging the decay time of the synaptic current (Pawelzik et al., 1999; Thomson et al., 2000; Kang-Park et al., 2004; Xu and Sastry, 2005; Gross et al., 2011). For instance, diazepam, a nonselective 1,4-BDZ, has been reported to potentiate both the amplitude and decay time of IPSCs in CA1 pyramidal neurons in hippocampal slices (Zhang et al., 1993). However, there are also examples of GABA_A_ PAMs such as stiripentol and CLB that prolong the decay of the synaptic current, with no effect on amplitude (Quilichini et al., 2006). Stiripentol selectively enhances the decay of IPSCs by increasing the open time of the receptor-channel (Quilichini et al., 2006) as does 1,4-BDZ derivative midazolam (Otis and Mody, 1992). GABA_A_ channels are also potentiated by barbiturates, which bind to a site independent of the benzodiazepine binding site and lengthen the mean open times with no effect on open probability (MacDonald et al., 1989). AZD7325 also prolonged the decay time of the currents without increasing the amplitude of IPSCs. This finding suggests that AZD7325 may prolong the mean open time of α_2_ / α_3_ containing GABA_A_ receptors similar to barbiturates or other AEDs that bind to the barbiturate site such as stiripentol, which also regulate IPSC decay kinetics without affecting the current amplitude (Zhang et al., 1993; Quilichini et al., 2006).

Prior work from our laboratory demonstrated that there is a strong strain dependence to the phenotype in the *Scn1a*^+/−^ mouse model of DS. Mice on a congenic 129S6/SvEvTac background are protected from seizures and premature mortality (Miller et al., 2014). Fine mapping of the Dravet syndrome modifier 1 (*Dsm1*) locus identified *Gabra2* as a potential candidate modifier gene that is differentially expressed between the seizure-resistant 129 strain and the seizure-susceptible F1 strain (Hawkins et al., 2016). Here we directly tested whether synapses known to contain α_2_ - GABA_A_ receptors are differentially modulated by AZD7325. The α subunits are known to govern the kinetics of GABA currents (Verdoorn et al., 1990; Lavoie et al., 1997), however we did not observe any difference in the amplitude or kinetics of mIPSCs between recordings from any of the mouse strains tested (not shown), likely because of the heterogeneous complement of receptors at synapses that contain both α_1_ and α_2_ receptors, where the α_1_ containing receptors govern the kinetics of IPSCs (Panzanelli et al., 2011). However, the effect of AZD7325 on mIPSC decays was significantly more pronounced in the seizure-resistant 129 strain than in the F1 and B6 seizure-susceptible strains, consistent with elevated abundance of α_2_ containing receptors at synapses in 129 strain relative to B6.

While we found a strain dependence of the effects of AZD7325 on mIPSCs, there was no genotype dependent effect in the DS mice. Synaptic currents were potentiated to a similar level in *Scn1a*^+/−^ mice as in wild-type littermates on the same congenic background, and this supports the notion that modifier genes in the background strains are important determinants of the phenotype expressed in DS mice. In addition, *in vivo* administration of AZD7325 to F1.*Scn1a*^+/−^ mice was associated with higher temperature threshold for hyperthermia-induced seizures in a dose dependent manner with no obvious sedative effect at the highest dose administered. This directly supports the therapeutic potential of α_2_ - selective PAMs as effective seizure suppressors. Prior studies have also suggested that α_2_-containing receptors contribute to seizure susceptibility. For instance, rats that are non-responders to typical AEDs after kindling have lower expression of several GABA_A_ receptor subunits including α_2_ in the hippocampus compared to kindled rats that are responsive to drug treatment (Bethmann et al., 2008). In addition, recent work demonstrated that loss of a novel α_2_ interacting partner, collybistin, led to lower α_2_ expression and loss of GABAergic synapses on the axon initial segment (AIS) along with greater seizure susceptibility (Hines et al., 2018). Furthermore, in human genetic studies a *de novo* mutation in the *GABRA2* gene that was found in patients with severe epilepsy and developmental delay, has been demonstrated to have lower channel expression and exhibit effects on gating of GABA_A_ receptors (Butler et al., 2018). Our studies in the *Scn1a*^+/−^ mouse model provide an explanation of how genetic modifiers can affect phenotype penetrance, and further support the importance of considering GABA receptor subunit selective pharmacology for the treatment and suppression of seizures.

## Acknowledgements

This work was supported by NIH grants R01 NS084959 (J.A.K.) and R01MH099114 (A.C.) and a seed grant from the American Epilepsy Society (A.C.). We thank Nick Brandon from AstraZeneca for providing AZD7325.

## Citations

Alhambra C et al. (2011) Development and SAR of functionally selective allosteric modulators of GABAA receptors. Bioorg Med Chem 19:2927–2938.

Bethmann K, Fritschy JM, Brandt C, Loscher W (2008) Antiepileptic drug resistant rats differ from drug responsive rats in GABA A receptor subunit expression in a model of temporal lobe epilepsy. Neurobiol Dis 31:169–187.

Brodie MJ (2017) Outcomes in newly diagnosed epilepsy in adolescents and adults: Insights across a generation in Scotland. Seizure 44:206–210.

Brunklaus A, Zuberi SM (2014) Dravet syndrome–from epileptic encephalopathy to channelopathy. Epilepsia 55:979–984.

Butler KM, Moody OA, Schuler E, Coryell J, Alexander JJ, Jenkins A, Escayg A (2018) De novo variants in GABRA2 and GABRA5 alter receptor function and contribute to early-onset epilepsy. Brain.

Calhoun JD, Hawkins NA, Zachwieja NJ, Kearney JA (2017) Cacna1g is a genetic modifier of epilepsy in a mouse model of Dravet syndrome. Epilepsia 58:e111–e115.

Fernandes HB, Riordan S, Nomura T, Remmers CL, Kraniotis S, Marshall JJ, Kukreja L, Vassar R, Contractor A (2015) Epac2 Mediates cAMP-Dependent Potentiation of Neurotransmission in the Hippocampus. J Neurosci 35:6544–6553.

Fritschy JM, Weinmann O, Wenzel A, Benke D (1998) Synapse-specific localization of NMDA and GABA(A) receptor subunits revealed by antigen-retrieval immunohistochemistry. J Comp Neurol 390:194–210.

Gingrich KJ, Roberts WA, Kass RS (1995) Dependence of the GABAA receptor gating kinetics on the alpha-subunit isoform: implications for structure-function relations and synaptic transmission. J Physiol 489 (Pt 2):529–543.

Greenfield LJ, Jr. (2013) Molecular mechanisms of antiseizure drug activity at GABAA receptors. Seizure 22:589–600.

Gross A, Sims RE, Swinny JD, Sieghart W, Bolam JP, Stanford IM (2011) Differential localization of GABA(A) receptor subunits in relation to rat striatopallidal and pallidopallidal synapses. Eur J Neurosci 33:868–878.

Gurrell R, Dua P, Feng G, Sudworth M, Whitlock M, Reynolds DS, Butt RP (2018) A randomised, placebo-controlled clinical trial with the alpha2/3/5 subunit selective GABAA positive allosteric modulator PF-06372865 in patients with chronic low back pain. Pain.

Hawkins NA, Kearney JA (2016) Hlf is a genetic modifier of epilepsy caused by voltage-gated sodium channel mutations. Epilepsy Res 119:20–23.

Hawkins NA, Zachwieja NJ, Miller AR, Anderson LL, Kearney JA (2016) Fine Mapping of a Dravet Syndrome Modifier Locus on Mouse Chromosome 5 and Candidate Gene Analysis by RNA-Seq. PLoS Genet 12:e1006398.

Hawkins NA, Anderson LL, Gertler TS, Laux L, George AL, Jr., Kearney JA (2017) Screening of conventional anticonvulsants in a genetic mouse model of epilepsy. Annals of clinical and translational neurology 4:326–339.

Hines RM, Maric HM, Hines DJ, Modgil A, Panzanelli P, Nakamura Y, Nathanson AJ, Cross A, Deeb T, Brandon NJ, Davies P, Fritschy JM, Schindelin H, Moss SJ (2018) Developmental seizures and mortality result from reducing GABAA receptor alpha2-subunit interaction with collybistin. Nat Commun 9:3130.

Jensen HS, Nichol K, Lee D, Ebert B (2014) Clobazam and its active metabolite N-desmethylclobazam display significantly greater affinities for alpha(2)-versus alpha(1)-GABA(A)-receptor complexes. PLoS One 9:e88456.

Jurgensen S, Castillo PE (2015) Selective Dysregulation of Hippocampal Inhibition in the Mouse Lacking Autism Candidate Gene CNTNAP2. J Neurosci 35:14681–14687.

Kang-Park MH, Wilson WA, Moore SD (2004) Differential actions of diazepam and zolpidem in basolateral and central amygdala nuclei. Neuropharmacology 46:1–9.

Kasugai Y, Swinny JD, Roberts JD, Dalezios Y, Fukazawa Y, Sieghart W, Shigemoto R, Somogyi P (2010) Quantitative localisation of synaptic and extrasynaptic GABAA receptor subunits on hippocampal pyramidal cells by freeze-fracture replica immunolabelling. Eur J Neurosci 32:1868–1888.

Kerti-Szigeti K, Nusser Z (2016) Similar GABAA receptor subunit composition in somatic and axon initial segment synapses of hippocampal pyramidal cells. Elife 5.

Klausberger T, Roberts JD, Somogyi P (2002) Cell type- and input-specific differences in the number and subtypes of synaptic GABA(A) receptors in the hippocampus. J Neurosci 22:2513–2521.

Knabl J, Witschi R, Hosl K, Reinold H, Zeilhofer UB, Ahmadi S, Brockhaus J, Sergejeva M, Hess A, Brune K, Fritschy JM, Rudolph U, Mohler H, Zeilhofer HU (2008) Reversal of pathological pain through specific spinal GABAA receptor subtypes. Nature 451:330–334.

Lavoie AM, Tingey JJ, Harrison NL, Pritchett DB, Twyman RE (1997) Activation and deactivation rates of recombinant GABA(A) receptor channels are dependent on alpha-subunit isoform. Biophys J 73:2518–2526.

Low K, Crestani F, Keist R, Benke D, Brunig I, Benson JA, Fritschy JM, Rulicke T, Bluethmann H, Mohler H, Rudolph U (2000) Molecular and neuronal substrate for the selective attenuation of anxiety. Science 290:131–134.

MacDonald RL, Rogers CJ, Twyman RE (1989) Barbiturate regulation of kinetic properties of the GABAA receptor channel of mouse spinal neurones in culture. J Physiol 417:483–500.

Manford M (2017) Recent advances in epilepsy. J Neurol 264:1811–1824.

McKernan RM et al. (2000) Sedative but not anxiolytic properties of benzodiazepines are mediated by the GABA(A) receptor alpha1 subtype. Nat Neurosci 3:587–592.

Miller AR, Hawkins NA, McCollom CE, Kearney JA (2014) Mapping genetic modifiers of survival in a mouse model of Dravet syndrome. Genes Brain Behav 13:163–172.

Mistry AM, Thompson CH, Miller AR, Vanoye CG, George AL, Jr., Kearney JA (2014) Strain- and age-dependent hippocampal neuron sodium currents correlate with epilepsy severity in Dravet syndrome mice. Neurobiol Dis 65:1–11.

Mody I, Pearce RA (2004) Diversity of inhibitory neurotransmission through GABA(A) receptors. Trends Neurosci 27:569–575.

Mullen SA, Scheffer IE (2009) Translational research in epilepsy genetics: sodium channels in man to interneuronopathy in mouse. Archives of neurology 66:21–26.

Mulligan MK, Wang X, Adler AL, Mozhui K, Lu L, Williams RW (2012) Complex control of GABA(A) receptor subunit mRNA expression: variation, covariation, and genetic regulation. PLoS One 7:e34586.

Nickolls SA et al. (2018) Pharmacology in translation: the preclinical and early clinical profile of the novel alpha2/3 functionally selective GABAA receptor positive allosteric modulator PF-06372865. Br J Pharmacol 175:708–725.

Nomura T, Oyamada Y, Fernandes HB, Remmers CL, Xu J, Meltzer HY, Contractor A (2016) Subchronic phencyclidine treatment in adult mice increases GABAergic transmission and LTP threshold in the hippocampus. Neuropharmacology 100:90–97.

Nusser Z, Sieghart W, Benke D, Fritschy JM, Somogyi P (1996) Differential synaptic localization of two major gamma-aminobutyric acid type A receptor alpha subunits on hippocampal pyramidal cells. Proc Natl Acad Sci U S A 93:11939–11944.

Nyiri G, Freund TF, Somogyi P (2001) Input-dependent synaptic targeting of alpha(2)-subunit-containing GABA(A) receptors in synapses of hippocampal pyramidal cells of the rat. Eur J Neurosci 13:428–442.

Olsen RW (2018) GABAA receptor: Positive and negative allosteric modulators. Neuropharmacology.

Olsen RW, Sieghart W (2008) International Union of Pharmacology. LXX. Subtypes of gamma-aminobutyric acid(A) receptors: classification on the basis of subunit composition, pharmacology, and function. Update. Pharmacol Rev 60:243–260.

Otis TS, Mody I (1992) Modulation of decay kinetics and frequency of GABAA receptor-mediated spontaneous inhibitory postsynaptic currents in hippocampal neurons. Neuroscience 49:13–32.

Panzanelli P, Gunn BG, Schlatter MC, Benke D, Tyagarajan SK, Scheiffele P, Belelli D, Lambert JJ, Rudolph U, Fritschy JM (2011) Distinct mechanisms regulate GABAA receptor and gephyrin clustering at perisomatic and axo-axonic synapses on CA1 pyramidal cells. J Physiol 589:4959–4980.

Pawelzik H, Bannister AP, Deuchars J, Ilia M, Thomson AM (1999) Modulation of bistratified cell IPSPs and basket cell IPSPs by pentobarbitone sodium, diazepam and Zn2+: dual recordings in slices of adult rat hippocampus. Eur J Neurosci 11:3552–3564.

Perucca P, Carter J, Vahle V, Gilliam FG (2009) Adverse antiepileptic drug effects: toward a clinically and neurobiologically relevant taxonomy. Neurology 72:1223–1229.

Prenosil GA, Schneider Gasser EM, Rudolph U, Keist R, Fritschy JM, Vogt KE (2006) Specific subtypes of GABAA receptors mediate phasic and tonic forms of inhibition in hippocampal pyramidal neurons. J Neurophysiol 96:846–857.

Quilichini PP, Chiron C, Ben-Ari Y, Gozlan H (2006) Stiripentol, a putative antiepileptic drug, enhances the duration of opening of GABA-A receptor channels. Epilepsia 47:704–716.

Ralvenius WT, Acuna MA, Benke D, Matthey A, Daali Y, Rudolph U, Desmeules J, Zeilhofer HU, Besson M (2016) The clobazam metabolite N-desmethyl clobazam is an alpha2 preferring benzodiazepine with an improved therapeutic window for antihyperalgesia. Neuropharmacology 109:366–375.

Rubinstein M, Westenbroek RE, Yu FH, Jones CJ, Scheuer T, Catterall WA (2015) Genetic background modulates impaired excitability of inhibitory neurons in a mouse model of Dravet syndrome. Neurobiol Dis 73:106–117.

Rudolph U, Knoflach F (2011) Beyond classical benzodiazepines: novel therapeutic potential of GABAA receptor subtypes. Nat Rev Drug Discov 10:685–697.

Saito A, Taniguchi Y, Rannals MD, Merfeld EB, Ballinger MD, Koga M, Ohtani Y, Gurley DA, Sedlak TW, Cross A, Moss SJ, Brandon NJ, Maher BJ, Kamiya A (2016) Early postnatal GABAA receptor modulation reverses deficits in neuronal maturation in a conditional neurodevelopmental mouse model of DISC1. Mol Psychiatry 21:1449–1459.

Scheffer IE, Berkovic S, Capovilla G, Connolly MB, French J, Guilhoto L, Hirsch E, Jain S, Mathern GW, Moshe SL, Nordli DR, Perucca E, Tomson T, Wiebe S, Zhang YH, Zuberi SM (2017) ILAE classification of the epilepsies: Position paper of the ILAE Commission for Classification and Terminology. Epilepsia 58:512–521.

Steel D, Symonds JD, Zuberi SM, Brunklaus A (2017) Dravet syndrome and its mimics: Beyond SCN1A. Epilepsia 58:1807–1816.

Taliani S, Cosimelli B, Da Settimo F, Marini AM, La Motta C, Simorini F, Salerno S, Novellino E, Greco G, Cosconati S, Marinelli L, Salvetti F, L’Abbate G, Trasciatti S, Montali M, Costa B, Martini C (2009) Identification of anxiolytic/nonsedative agents among indol-3-ylglyoxylamides acting as functionally selective agonists at the gamma-aminobutyric acid-A (GABAA) alpha2 benzodiazepine receptor. J Med Chem 52:3723–3734.

Thomson AM, Bannister AP, Hughes DI, Pawelzik H (2000) Differential sensitivity to Zolpidem of IPSPs activated by morphologically identified CA1 interneurons in slices of rat hippocampus. Eur J Neurosci 12:425–436.

Verdoorn TA, Draguhn A, Ymer S, Seeburg PH, Sakmann B (1990) Functional properties of recombinant rat GABAA receptors depend upon subunit composition. Neuron 4:919–928.

Whiting PJ (2003) GABA-A receptor subtypes in the brain: a paradigm for CNS drug discovery? Drug Discov Today 8:445–450.

Wirrell EC, Laux L, Donner E, Jette N, Knupp K, Meskis MA, Miller I, Sullivan J, Welborn M, Berg AT (2017) Optimizing the Diagnosis and Management of Dravet Syndrome: Recommendations From a North American Consensus Panel. Pediatric neurology 68:18–34.e13.

Xu JY, Sastry BR (2005) Benzodiazepine involvement in LTP of the GABA-ergic IPSC in rat hippocampal CA1 neurons. Brain Res 1062:134–143.

Xu-Friedman MA, Regehr WG (2000) Probing fundamental aspects of synaptic transmission with strontium. J Neurosci 20:4414–4422.

Yeo S, Hodgkinson CA, Zhou Z, Jung J, Leung M, Yuan Q, Goldman D (2016) The abundance of cis-acting loci leading to differential allele expression in F1 mice and their relationship to loci harboring genes affecting complex traits. BMC genomics 17:620.

Yu FH, Mantegazza M, Westenbroek RE, Robbins CA, Kalume F, Burton KA, Spain WJ, McKnight GS, Scheuer T, Catterall WA (2006) Reduced sodium current in GABAergic interneurons in a mouse model of severe myoclonic epilepsy in infancy. Nat Neurosci 9:1142–1149.

Zhang L, Weiner JL, Carlen PL (1993) Potentiation of gamma-aminobutyric acid type A receptor-mediated synaptic currents by pentobarbital and diazepam in immature hippocampal CA1 neurons. J Pharmacol Exp Ther 266:1227–1235.

Zhou D, Sunzel M, Ribadeneira MD, Smith MA, Desai D, Lin J, Grimm SW (2012) A clinical study to assess CYP1A2 and CYP3A4 induction by AZD7325, a selective GABA(A) receptor modulator - an in vitro and in vivo comparison. Br J Clin Pharmacol 74:98–108.

